# The role of rat prelimbic cortex in decision making

**DOI:** 10.1101/2024.03.18.585593

**Authors:** Jensen A. Palmer, Samantha R. White, Kevin Chavez Lopez, Mark Laubach

## Abstract

The frontal cortex plays a critical role in decision-making. One specific frontal area, the anterior cingulate cortex, has been identified as crucial for setting a threshold for how much evidence is needed before a choice is made (Domenech & Dreher, 2010). Threshold is a key concept in drift diffusion models, a popular framework used to understand decision-making processes. Here, we investigated the role of the prelimbic cortex, part of the rodent cingulate cortex, in decision making. Male and female rats learned to choose between stimuli associated with high and low value rewards. Females learned faster, were more selective in their responses, and integrated information about the stimuli more quickly. By contrast, males learned more slowly and showed a decrease in their decision thresholds during choice learning. Inactivating the prelimbic cortex in female and male rats sped up decision making without affecting choice accuracy. Drift diffusion modeling found selective effects of prelimbic cortex inactivation on the decision threshold, which was reduced with increasing doses of the GABA-A agonist muscimol. Stimulating the prelimbic cortex through mu opioid receptors slowed the animals’ choice latencies and increased the decision threshold. These findings provide the first causal evidence that the prelimbic cortex directly influences decision processes. Additionally, they suggest possible sex-based differences in early choice learning.

---

The frontal cortex has been well established across species as being involved in decision making processes, with numerous studies reporting signals from this region linked to decision making (Hanks and Summerfield, 2017). Drift Diffusion Modeling (Ratcliff et al., 2016) is commonly used to better understand the neural signals and computations involved in two-choice decision making tasks. Notably, a recent fMRI study found evidence for activity in the anterior cingulate cortex (ACC) tracking the decision threshold (Domenech and Dreher, 2010). Threshold accounts for the amount of information needed to make a choice, and is one of the two main measures provided by drift diffusion models. The other is the drift rate, how quickly information is integrated into a choice.

In rodents, the prelimbic cortex (PLC) is a rostral component of the ACC, and may be homologous to the pregenual ACC in primates (Laubach et al., 2018). The PLC has been shown to play a role in instrumental learning (Corbit and Balleine, 2003; Killcross and Coutureau, 2003; Tran-Tu-Yen et al., 2009) and inhibitory control (Chudasama and Muir, 2001; Risterucci et al., 2003; Narayanan et al., 2006). To our knowledge, no study has examined the role of the rodent ACC in decision making with regards to the parameters of drift diffusion modeling. The major goal of this study was to assess the role of the rodent frontal cortex in decision making, specifically if the PLC in rodents is similarly involved in maintaining decision threshold, as Domenech and Dreher (2010) found in humans.

To study the role of PLC in decision making, we used a behavioral design developed by White et al. (2024) to selectively study choice learning. A standard two-alternative forced-choice design was used with open-source LED matrices as visual stimuli (Swanson et al., 2021). White et al. (2024) used a unique training method to separate learning the values of task stimuli from learning to make choices between the stimuli. They trained male rats to learn the reward values of stimuli by presenting single stimuli on each trial, over a series of “value learning” sessions. Then, they tested the rats with choices between the stimuli, over a series of “choice learning” sessions. They found that male rats responded more slowly on choice trials compared to trials with single offers and came to need less information needed to make a choice during early choice learning.

Here, we used the same behavioral task to investigate the role of the PLC in decision making. Intra-cortical pharmacological methods were used to reversibly inactivate PLC by infusing the GABA-A receptor agonist muscimol and stimulate the cortex by infusing the mu opioid agonist DAMGO. Stimulation of mu opioid receptors disinhibits the PLC through actions on parvalbumin interneurons (Lau et al., 2020). Inactivating PLC sped up performance in both males and females without affecting choice accuracy. By contrast, stimulating the PLC through mu-opioid receptor stimulation had the opposite effects, slowing performance again without affecting choice accuracy. To understand how these perturbations of PLC affected the decision process, we used hierarchical Bayesian drift diffusion modeling (Wiecki et al., 2013). We found that inactivation of PLC selectively reduced the decision threshold, without affecting the drift rate. Exciting the PLC slowed performance, increased the decision threshold, and also reduced the drift rate in both males and females.

Prior to carrying out these experiments, we noted differences in how female and male rats learned the task. We observed that males, but not females, became faster at responding during early learning and females responded faster and more selectively to the stimulus that yielded a higher value reward compared to male rats. To examine if these differences were associated with differences in decision processing by the female and male rats, we fit drift diffusion models and found that decision threshold decreased in males, but not females, over the initial period of choice learning. Drift rate was unaffected by learning, but females had greater drift rates than males across sessions. We further noted differences in the shapes of the response time distributions in the female and male rats. To examine these differences, we used ExGauss models (Heathcote et al., 1991) to decompose the response time distributions into components that account for mean and variance of the distributions. These analyses revealed a clear difference in how female and male rats responded on trials with single and dual offers. Females showed shorter mean latencies with higher variability compared to males. These findings might reflect a higher level of exponential variability in the decision process of the female rats (“decision noise”: Holhe, 1965). Alternatively, the higher level of exponential could reflect value processing, as variability of response times by the females was highest when they were forced to respond to the low-value stimuli on single offer trials. Together, our studies demonstrate a causal role of the prelimbic cortex in the decision process and suggest a potential sex difference in how rats learn two-alternative forced-choice tasks.

## Materials and Methods

Procedures were approved by the Animal Care and Use Committee at American University (Washington, DC) and conformed to the standards of the National Institutes of Health Guide for the Care and Use of Laboratory Animals. Data from these studies are available on GitHub: https://github.com/LaubachLab/PLC-DDM

### Animals

Nine male Long Evans rats (350-450g, five from Charles-River, four from Envigo) and nine female Long Evans rats (200-250g, Envigo) were used in this study. Animals were individually housed on a 12/12 h light/dark cycle. During training and testing, animals had regulated access to food (12-16 grams) to maintain body weights at ∼90% of their free-access weights. Two males were not motivated by liquid sucrose rewards, so they were switched to water regulation early in training. They were maintained at ∼90% of their free-access weights. These animals typically consumed 10-15 ml of water during behavioral sessions and were given an additional 10-15 ml around 4 PM daily with free access to food. They had one day per week of free access to water.

### Behavioral apparatus

Animals were trained in sound-attenuating behavioral boxes (Med Associates) that had a single, horizontally placed spout mounted to a lickometer 6.5 cm from the floor with a single white LED placed 4 cm above the spout. Solution lines were connected to 60cc syringes and solution was made available to animals by lick-triggered, single speed pumps (PHM-100; Med Associates) which drove syringe plungers. Each lick activated a pump which delivered roughly 30 μL per 0.5 second activation. On the wall opposite the spout, three 3D-printed nosepoke ports were aligned 5 cm from the floor and 4 cm apart, and contained Adafruit 3-mm IR Break Beam sensors. A Pure Green 1.2” 8×8 LED matrix (Adafruit) was placed 2.5 cm above the center of each of the three nose poke ports outside of the box for visual stimulus presentation. LED matrices are controlled using Adafruit_GFX and Adafruit_LEDBackpack libraries, and microcontroller software is provided in a previous publication (Swanson et al., 2021).

### Training procedure: Value Learning

Nine males and nine females were trained in a behavioral task previously described in White et al. (2024) (Figure 1). First, animals licked at a reward spout in an operant chamber to receive 16% wt/vol liquid sucrose with the LED above the spout turned on. In subsequent sessions, animals were trained using the method of successive approximations to respond in nose poke ports in response to distinct visual stimuli. A 4x4 square of illuminated LEDs displayed over the center port signaled animals to initiate trials. Trial initiation was followed by lateralized presentation of one of two cues, which will be referred to as “single-offer” trials (Figure 1B). The cues were high-luminance (eight illuminated LEDs) or low-luminance (two illuminated LEDs), and were presented randomly by side (left or right) and luminance intensity. The location of illuminated LEDs changed every millisecond during the period of illumination, which began when rats entered the central port and ended when rats entered one of the lateralized ports. Responses for high-luminance yielded access to 16% wt/vol sucrose reward at the reward spout, and low-luminance yielded a 4% wt/volume sucrose reward. Responses to non-illuminated ports were considered as errors and were unrewarded. Responses that took longer than 5 seconds following trial initiation were counted as errors of omission and were unrewarded. On valid trials, animals had to collect their reward within 5 seconds following responses to receive fluid. Animals were trained for five, 60-minute sessions of at least 100 trials in single-offer sessions before moving on to dual-offer acquisition and test sessions.

**Figure 1.**
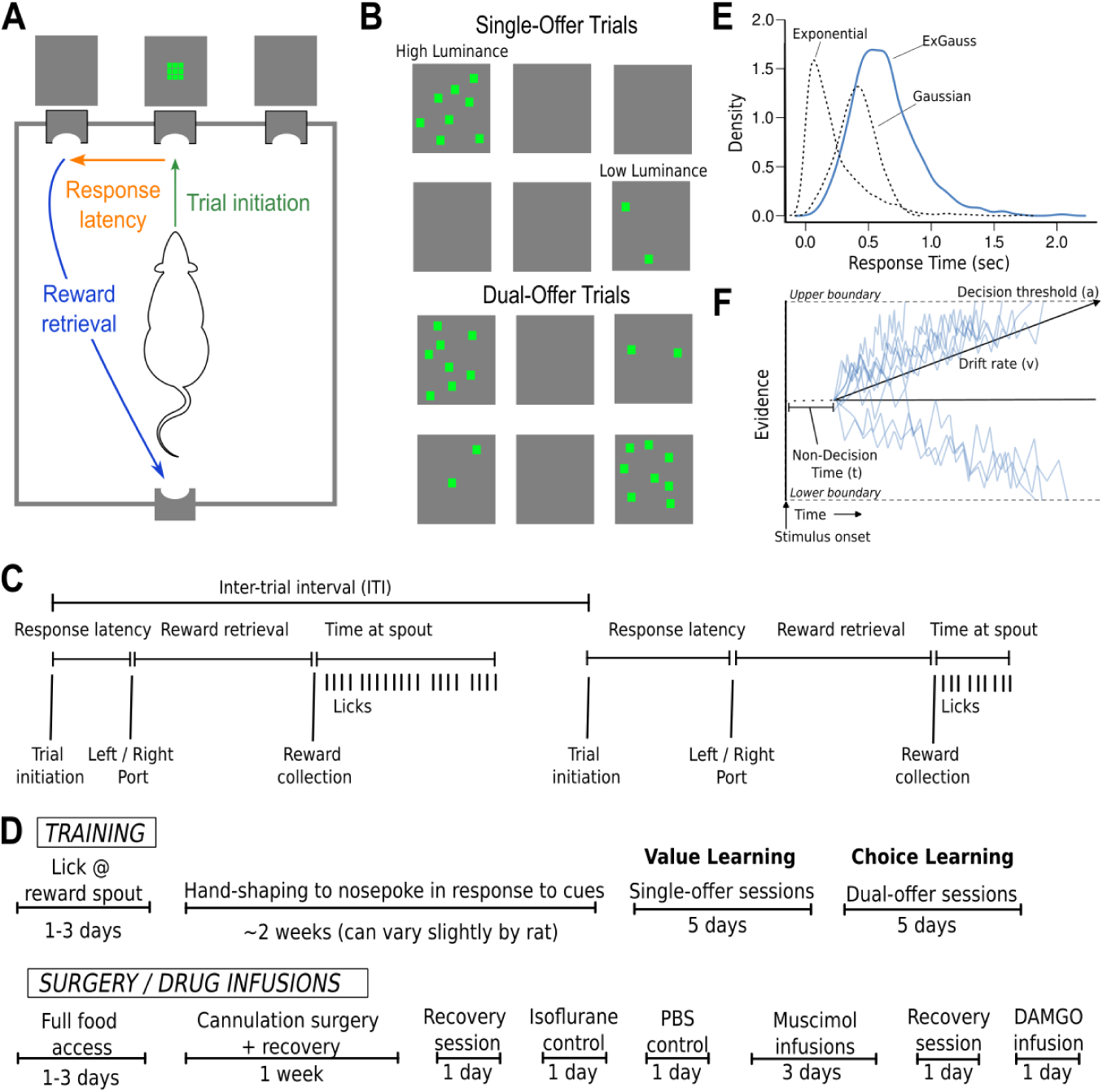
Experimental design and data analysis. A. Sequence of task events. Rats initiated trials by entering a central port. This extinguished a visual stimulus presented above the port. Then, a stimulus was presented above one or both of the side ports. Rats moved from the center port to one of the side ports to choose the stimulus. The time taken to respond in the side ports was the response latency. The rats were then required to contact a spout, located inside a reward port on the opposite side of the behavioral chamber. The time taken to travel from the side ports to the reward port was the reward retrieval latency. B. Stimulus set as they appear on LED grids. Single-offer trials (top) may have presentation of high-luminance or low-luminance cues. Dual-offer trials (bottom) have high-luminance and low-luminance cues presented simultaneously. LEDs were activated for 1 ms at a randomly selected location on the LED grids and the patterns of activated LEDs fluctuated over locations throughout the period of stimulus presentation. Stimuli terminated when rats entered one of the side ports. C. Sequence of behavioral events over two example trials. D. Timeline of training, surgery, and drug infusions. E. Schematic of the ExGauss distribution and its parameters. The ExGauss distribution is the sum of a Gaussian distribution and an exponential distribution. The Gaussian distribution accounts for the peak in the distribution of response latencies. The exponential distribution accounts for the positive tail of the response latency distribution. F. Drift diffusion model and its parameters. Blue lines represent the evidence for choosing between two options. The decision process begins after the stimulus has been encoded (part of what is called the non-decision time). Evidence accumulates towards one of two decision boundaries, i.e., choose the brighter or dimmer stimulus. The rate at which evidence accumulates is accounted for by the drift rate. The difference between the boundaries is the decision threshold, i.e., how much evidence is needed to trigger a choice.

### Training procedure: Choice Learning

As in training, animals initiated trials by nose poking at the center port. In 60-minute sessions, two-thirds (∼67%) of trials were single-offer trials, as described above. In the remaining third (∼33% of trials), animals were presented with both the high- and low-luminance cues simultaneously, randomized by side, and the animals had the choice of responding to either cue. These trials are referred to as “dual-offer” trials (Figure 1B). Single- and dual-offer trials were interleaved throughout the 60-minute sessions. The rats always received 16% sucrose at the reward port after they responded in the port below the high-luminance cue and always received 4% sucrose after they responded in the port below the low-luminance cue, and errors are defined in the same way as described for the value learning stage of the training. Animals experienced choice learning over five 60-minute sessions before undergoing surgery. A difference in choice learning between the present study and White et al. (2024) is that the animals only experienced choice sessions during this stage of training. In the study by White et al. (2024), there were sessions with only single offer trials interleaved between sessions with choice learning. All test sessions following surgery included mixtures of single-offer and dual-offer trials, as described here for the choice learning phase of the task.

The behavioral measures of interest are referred to as “latency,” “reward retrieval,” “time at spout,” “inter-trial interval,” “high value preference,” and “side bias,” (Figure 1A,C). Latency is measured as the time taken for rats to go from trial initiation to responding in an illuminated choice port after the onset of the visual stimuli. Reward retrieval is measured as the time it takes from making a nose poke response to the time that reward is collected. Only valid trials with latencies and reward retrievals that were less than five seconds were included in analyses. Time at spout is measured as the time between the first lick when a reward is collected and the last lick before a new trial is initiated. Inter-trial interval (ITI) is measured as the time from one trial initiation until the next trial initiation. Only trials where ITI is less than 60 seconds were included in analyses. High value preference is measured as the ratio of dual-offer trials that the animals responded to the high-luminance cue. Side bias is measured as the percentage difference from 50% that a rat chooses a given side on dual-offer trials. For example, if a rat chooses left 70% and right 30% of dual-offer trials, then this rat would have a 20% side bias.

### Surgery

Animals were given two to three days with free access to food and water prior to cannula implantation surgery. Anesthesia was induced by intraperitoneal injection of ketamine (100 mg/kg for males, 75 mg/kg for females) and maintained with isoflurane (0.5-2.0%; flow rate 5.0 cc/min). A bolus of carprofen was also intraperitoneally injected in the animals immediately following the ketamine injection. Animals were placed into a stereotaxic frame using non-penetrating ear bars. The scalp was then shaved and covered with iodine, and the eyes were covered with ophthalmic ointment. Lidocaine (0.5 mL) was injected under the scalp and a longitudinal incision was made along the skull, followed by lateral retraction of the skin. Two skull screws were placed on the caudal edges of the skull to support adhesion of the implant. Bilateral craniotomies were made above implant locations targeting the prelimbic cortex. Twenty-six gauge stainless steel guide cannula (Plastics One) were lowered into the rostral PLC (coordinates from bregma AP +3.0mm, ML +/-1.2mm, DV -2.2mm surface of the brain at a 30-degree lateral and 12-degree posterior angle) (Paxinos and Watson, 2014). Coordinates were adjusted to avoid blood vessels, leadings to most placements being anterior to the starting coordinates. The guide cannula contained 33 gauge stainless steel wire that projected 0.4mm past the tip of the guide cannula. Craniotomies were closed using cyanoacrylate (Slo-Zap) and an accelerator (Zip Kicker), and methyl methacrylate dental cement (AM Systems) was applied around the implants and affixed to the skull via the skull screws. Animals were given 0.5 mL of carprofen in 500 mL of water for postoperative analgesia and recovered in their home cages for one week with full food and water with daily monitoring until weights returned to presurgical levels. Animals returned to regulated food or water access before continuing with behavioral testing.

### Drug infusions

The timeline of drug infusions is summarized in Figure 1D. Following recovery from surgery and regulation of food or water access, animals were first re-acclimated to the behavioral task until they performed as prior to surgery. Then, animals were exposed to the same duration and levels of isoflurane gas used during drug infusions to control for exposure to isoflurane. Drugs were infused under isoflurane gas to limit effects of experimenter handling, as in previous studies from our lab (e.g. Narayanan et al., 2006; White and Laubach, 2022). A second control was carried out the following day where animals received an infusion of the same volume (1 μl) of PBS without drug into the rostral PLC while anesthetized. The following day a series of three muscimol infusions of increasing concentrations taking place over three consecutive days began (0.01 μg/μl, 0.1 μg/μl, 1.0 μg/μl). Following muscimol test sessions, animals were run in a test session with no infusions to serve as a local control for an infusion of DAMGO ([d-Ala2, N-Me-Phe4, Gly5-ol]- enkephalin) (1.0 μg/μl; Giacomini et al., 2021 and White and Laubach, 2022), which took place the following day. A single dose of DAMGO was used due to the repeated drug infusions that were done during the inactivation sessions. At least one week was allowed for washout of DAMGO. Eight animals (four males and four females) were subsequently tested with unilateral infusions of muscimol (1.0 μg/μl) over two consecutive days, counterbalancing which hemisphere received infusions first. These additional infusions were done to test for effects of inactivation on side bias.

All drugs were obtained from Tocris and made into solutions using sterile PBS (pH 7.4). Infusions occurred by inserting a 33-gauge injector into the guide cannula that was flush with the tip of the guide cannula. The injector was connected to a 10μL Hamilton syringe via 0.38 mm diameter polyethylene tubing. A volume of 1.0 μL of fluid was delivered at a rate of 0.25 μL/min with a syringe infusion pump (KDS Scientific). The injector was left in place for two minutes after completion of the infusion to allow for diffusion of the solution, after which the injector was removed and the dummy cannula was replaced. Animals were tested in dual-offer sessions one hour after muscimol infusions and 30 minutes after DAMGO infusions.

### Confirmation of cannula placement

Following all experimental test sessions, animals were anesthetized with isoflurane and injected intraperitoneally with Euthasol. Animals were then transcardially perfused with 500 ml of saline solution followed by 500 ml of 4% paraformaldehyde. Brains were removed and post-fixed in 4% paraformaldehyde overnight and were then transferred to solutions containing 20% sucrose and 20% glycerol. Brains were sliced into 60 μm coronal sections using a freezing microtome and mounted onto gelatin-coated slides for Nissl staining via thionin. The thionin-treated sections were dried through a series of alcohol steps, covered with Clearium, and coverslipped. Sections were imaged using a Tritech Research scope (BX-51-F), Moticam Pro 282B camera, and Motic Images Plus 2.0 software. The most ventral point of the cannula track was compared against the Paxinos and Watson (2014) atlas to confirm coordinates of infusion sites. That version of the atlas uses the term Area 32 to indicate the region of frontal cortex that is also called prelimbic cortex.

### Statistical analysis

Behavioral events were recorded through MED-PC and extracted through custom scripts written in Python (Anaconda distribution: https://www.continuum.io/). All statistical analyses were carried out using R (R Core Team 2023) run in Jupyter notebooks (http://jupyter.org/) via rpy2. Statistical tests were performed using repeated measures ANOVA’s (R: “aov”) to account for within-subject effects of session (i.e. training session or drug infusion), sex, trial type (single-offer versus dual-offer), value (high versus low stimulus/reward), and the interactions between these variables on measures of responding (i.e. latency, reward retrieval, time at spout, ITI). Dependent variables were logarithmically transformed. Reported statistics include p-values and F-statistics. Where applicable, Tukey post hoc testing (R: “TukeyHSD”) was used to report significant pairwise differences. Spearman rank correlation coefficients were used to compare side bias to high value preference (R: “cor.test”). Each rat was dropped from statistical tests to confirm that effects were observed in all rats and that there were no differences between animals motivated by regulated access to food and the two male rats motivated by regulated access to water.

### ExGauss modeling

Median response latencies were used as the primary measure of the speed of performance in this study. ExGauss modeling was used to further understand how learning and perturbation of the prelimbic cortex influenced performance. It is a data analysis approach that approximates the distribution of response times as arising from the combination of two statistical processes (Figure 1E). A Gaussian component accounts for the mean response time. An exponential component accounts for the “long tail” variance of the response times. Together, the two components account for the overall shape of the response time distribution (see Luce, 1986, Heathcote et al., 1991, and Matzke and Wagenmakers, 2009 for review).

ExGauss models have three parameters. They are mu, the mean of the Gaussian distribution, sigma, the standard deviation of the Gaussian distribution, and tau, the exponential distribution. All three parameters were estimated, but the sigma parameter was not found to vary in any systematic fashion, and was not reported below. As there were few low value choices on dual-offer trials compared to the other trial types, we focused on differences between high and low value single-offer trials and high value dual-offer trials in interpreting results from the ExGauss models.

To estimate the parameters of the ExGauss models, we used the MMest.ExGauss function from the RobustEZ package (Wagenmakers et al. 2008). Effects of the ExGauss parameters on behavior was first assess using MANOVA (statsmodels library for Python). rmANOVA was then used to determine effects of session, sex, trial type, value, and the interactions between these variables on the ExGauss parameters.

### Drift Diffusion modeling

The HDDM package (Wiecki et al., 2013; version 0.9.6) was used to quantify effects of learning, sex, and drugs on decision making. We used the HDDM package to estimate three key parameters of DDM models, the drift rate, decision threshold, and non-decision time (Figure 1F). Drift rate accounts for how quickly the rats integrate information about the stimuli. Threshold accounts for how much information is needed to trigger a decision. Non-decision time accounts for the time taken to initiate stimulus integration and execute the motor response (choice). A fourth parameter that can be included in HDDM models is called bias (not shown in Figure 1F). It accounts for variability in the starting point of evidence accumulation. We used a fixed bias of 0.5 for the main analyses reported in this study.

HDDM models were fit that allowed for a single DDM parameter (drift rate, threshold, non-decision time) to vary freely over sessions (choice learning or drug dose). The other parameters were estimated globally (i.e., using data from all sessions in a given experiment: learning, inactivation, or DAMGO). Models were trained using data from dual-offer trials in all sessions and animals, and not for single sessions or animals. HDDM models were fit by running version 0.9.6 of the package under Python 3.7. Parameters were from Pedersen et al. (2021): Models were run five times, each with 5000 samples and the first 2500 samples were discarded as burn-in. A fixed bias term of 0.5 was used. Convergence was validated based on the Gelman-Rubin statistic (Gelman and Rubin, 1992). The autocorrelations and distributions of the parameters for each parameter and predictions of the response latency distributions for each animal were visually assessed to further assess convergence. HDDM is sensitive to outliers (Wiecki et al., 2013), so we included latencies up to the 95th percentile of the distribution in our analyses. Exploratory data analysis found that the 95th percentile cutoff was approximately the same for the response time distributions in males and females.

We fit an additional set of HDDM models in which we allowed the bias term to freely vary over rats and using global estimates of the three main parameters of the DDM model. The deviance information criterion for models with and without the bias parameter were approximately equal, indicating that the fits of models with and without the bias term were generally the same. The bias parameter was estimated as being only slightly offset from 0.5 (mean 0.5153, standard deviation 0.025) towards the bounds for the higher value option. These results were obtained from models with bias freely varying over rats and other parameters estimated globally. There were more noticeable fluctuations in traces of the posteriors and higher levels of autocorrelation for the three main DDM parameters in models in which one of the parameters freely varied over rats and bias was estimated globally. For this reason, we report results from HDDM models with bias fixed at 0.5 in the results reported below.

## Results

### Effects of early learning on performance and decision making

Over the five sessions of value learning, all animals reduced their median response latencies (p=1.06e-06, F(4,142)=9.295) (Figure 2A). Latencies were consistently shorter when rats responded to the high value stimulus compared to the low value stimulus (p=1.83e-15, F(1,142)=80.005) (Figure 2A). Females had shorter latencies than males across all training sessions (p=0.0422, F(1,15)=4.930) (Figure 2A). Reward retrieval (p=1.03e-05, F(4,139)=7.816) and ITI (p=3.91e-05, F(4,140)=6.952) also decreased over single-offer sessions, and females retrieved rewards more quickly than males (p=0.01, F(1,15)=8.852) (not shown).

**Figure 2.**
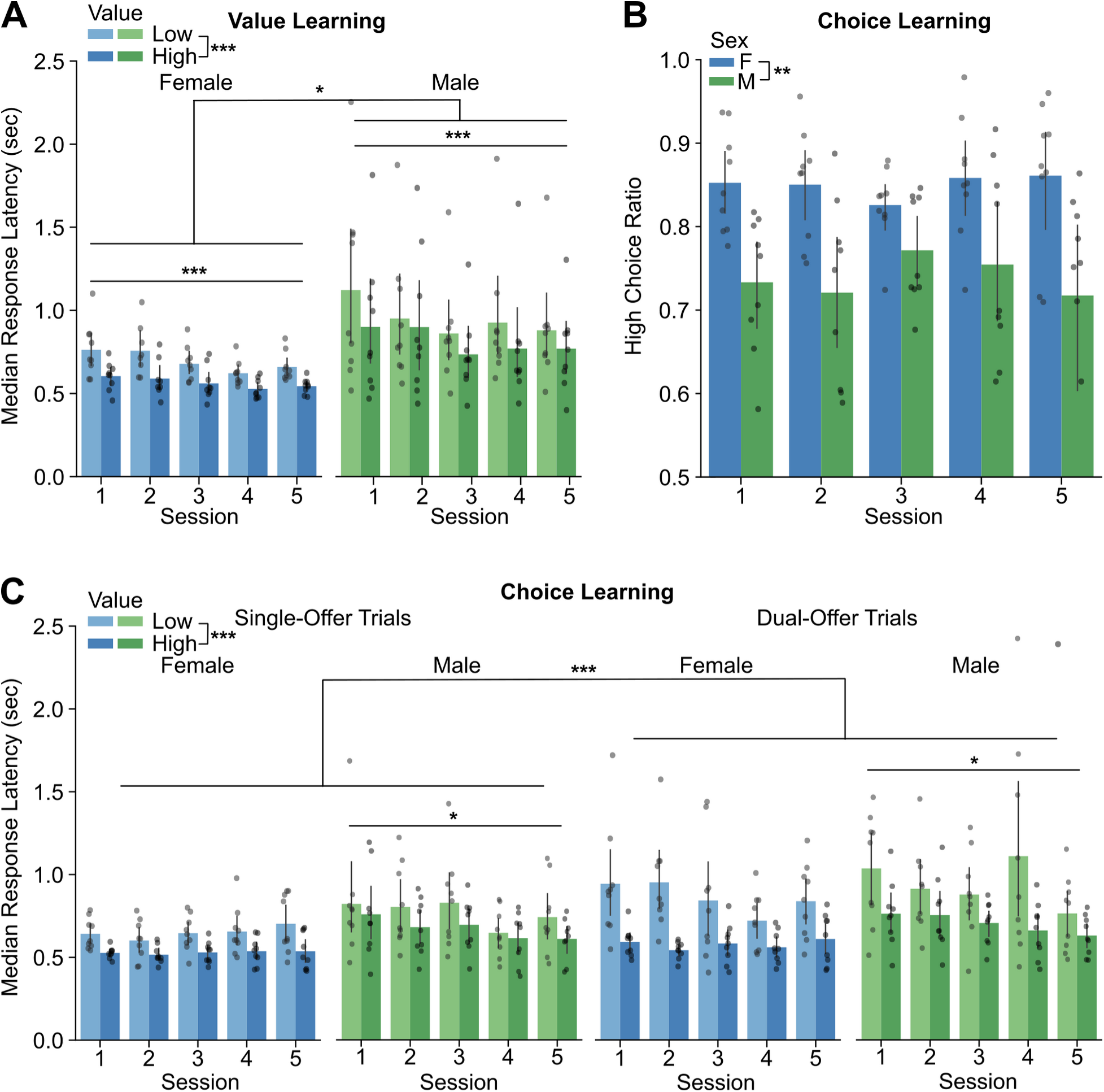
Effects of choice learning on preference and latency. A. Median response latencies for the high versus low-luminance stimulus over value learning sessions, broken up by sex. Latencies were briefer in females compared to males. Latencies decreased over the sessions with value learning. B. The preference for the high-value stimulus was greater in females compared to males over the period of choice learning. C. Median response latencies for the high versus low-luminance stimulus over choice learning sessions, broken up by sex and trial type (number of offers). As during value learning, females responded more quickly than males. Response latencies decreased over the sessions with choice learning, especially for when the rats chose the high-value option. Latencies were briefer for trials with choices of high-value stimuli, compared to low-value stimuli, over all stages of learning. Asterisks: *** p<0.001, ** p<0.01, * p<0.05.

Over the next five sessions of choice learning, the rats experienced dual-offer trials on one-third of the trials. Females demonstrated greater high-value preference than males across all five sessions (p=0.00103, F(1,16)=15.99) (Figure 2B). Latencies to the high-value stimuli were briefer for all rats compared to latencies to the low value stimulus (p=< 2e-16, F(1,304)=128.948) (Figure 2C). Latencies were longer for dual offer trials compared to single-offer trials (p=7.59e-11, F(1,304)=45.534) (Figure 2C). There was a significant interaction between session and sex (p=0.03027 F(4,304)=2.711) that revealed that males, but not females, reduced their latencies over sessions (Figure 2C). A significant interaction between value and sex (p= 0.02255, F(1,304)=5.257) was also found, with females responding more quickly than males specifically to the high value stimulus.

Beyond these changes in response latency, we also noted effects of learning on reward retrieval. There was a significant interaction between session and sex (p=0.00905, F(4,304)=3.441), due to a decrease in retrieval times over sessions only in males. All animals spent more time at the reward spout when a high value reward was consumed (p=<2e-16, F(1,300)=183.612) and did not show any effects of side bias during learning (p=0.822, F(4,64)=0.38).

To understand how choice learning affected choice latency in more detail, we used ExGauss modeling (Heathcote et al., 1991) to estimate effects of training session, sex, stimulus value, and the number of offers on the shape of the latency distributions. The Gaussian means (mu) of the response latency distributions were lower overall for females compared to males (p=0.00104, F(1,14)=16.997) (Figure 3A). There was also a significant effect of the number of offers on mu (p=0.00966, F(2,218)=4.74). Post hoc testing revealed that mu was significantly lower for single-offer, high value trials than single-offer, low value trials (p=0.0338613) (Figure 3A). The other main measure from ExGauss modeling (tau) accounts for exponential variability in the response times and the long tail of the response time distribution. Males had lower overall tau (p=0.0433, F(1,14)=4.938) and showed a reduction in tau over sessions with choice learning (p=5.15e-05, F(4,218)=6.577) (Figure 3B). A significant interaction between the number of offers and sex was found (p=6.6e-06, F(2,109)=13.334). Post hoc testing revealed that in females, tau was significantly greater in single-offer, low value trials compared to both single-offer, high value (p=0.0000064) and dual-offer, high value (p=0.0062493) trials. (Figure 3B).

**Figure 3.**
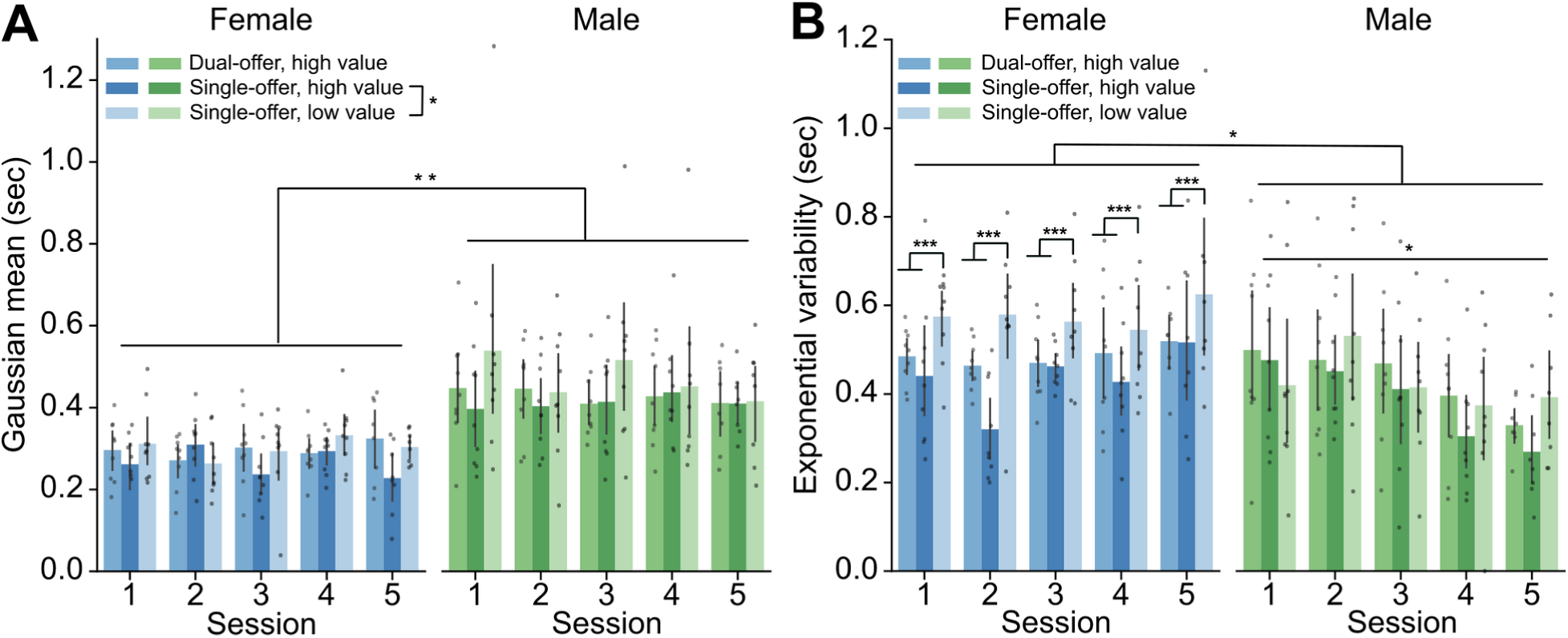
ExGauss modeling of response latencies during choice learning. This analysis decomposed response time variability into two components. One, called mu, accounts for the peak in the response time distributions, reflective of the Gaussian mean. The other, called tau, accounts for the long tail in the response time distributions. Comparisons of these parameters were made for dual-offer high value trials, single-offer high value trials, and single-offer low value trials. **A.** Mu was lower in females compared to males across all sessions of choice learning. Mu was also lower across trial types for responses to single offers of the high-value stimulus compared to the low-value stimulus. **B.** Tau was higher in females compared to males across all sessions of choice learning. Tau was also higher on trials with single offers of the low-value stimulus compared to single and dual offer trials with the high-value stimulus. Finally, tau reduced over sessions with choice learning on dual offer trials in males, but not females. Asterisks: *** p<0.001, ** p<0.01, * p<0.05.

To understand how choice learning affected the decision process, we used Drift Diffusion modeling using the HDDM package (Wiecki et al., 2013). These models use a hierarchical Bayesian approach, in which thousands of models are generated and blended over the collection of rats and sessions. As a result, traditional statistics, such as ANOVA, cannot be used to assess results from HDDM models. Therefore, we used an approach that is standard in the literature for reporting results from HDDM models, based on Bayesian confidence intervals for the parameters from the models. Significant effects based on these measures are denoted by asterisks in Figure 4.

**Figure 4.**
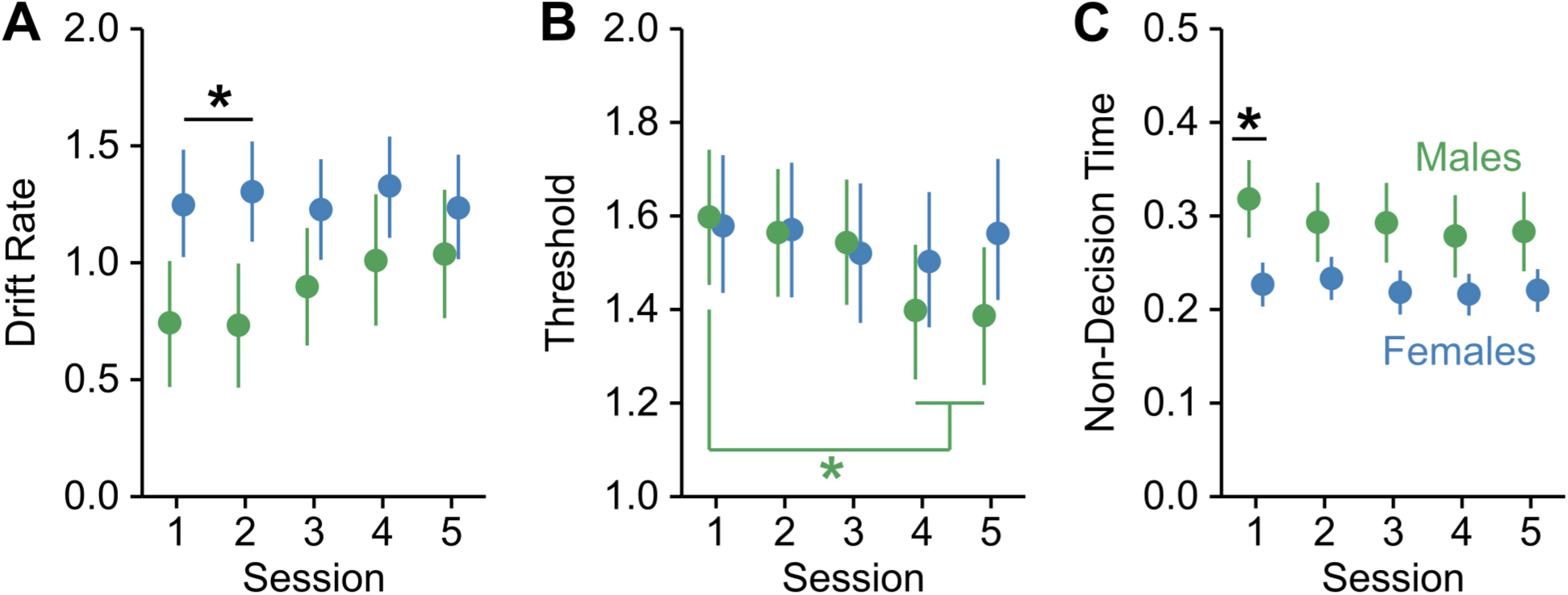
Drift diffusion modeling during choice learning. A. Drift rate did not change with choice learning. However, females had higher drift rates than males in the first two sessions with choice learning. B. Threshold decreased with choice learning in males, but not females. Threshold in the first session with choice learning was higher compared to the fourth and fifth sessions in males. C. Non-decision time did not change with learning. However, males had longer non-decision times compared to females in the first session of choice learning. Asterisks: *** p<0.001, ** p<0.01, * p<0.05.

To assess effects of learning in the female and male rats, we fit separate HDDM models to the groups of male and female rats. We noted differences over sessions in the drift rate, threshold, and non-decision time parameters based on the posterior distributions of the estimated parameter values. Neither males nor females showed any effects of learning on the posterior distributions of drift rate or non-decision time (Figure 4A,C). Models based on data from the male, but not the female, rats showed a decrease in the mean value of the threshold (Figure 4B). For males, the probability that threshold was larger in the first session with choice learning than in the fourth and fifth learning sessions was 0.0228 and 0.0256, respectively. For females, the probability that threshold was larger in the first session compared to the fifth session was 0.442. The 95% confidence intervals for drift rate from the male and female rats did not overlap in the first two sessions with choice learning, indicating that the females had higher drift rates compared to males early in choice learning. The 95% confidence intervals for non-decision time from the female and male rats did not overlap in the first session of choice learning, indicating that the males took more time for non-decision processes in the first session of choice learning.

### Effects of reversible inactivation of the prelimbic cortex

To understand the role of the prelimbic cortex (PLC) in decision making, we implanted the animals with infusion cannula (Figure 5) and tested them in the choice version of the task after infusions of increasing doses of muscimol. Reversible inactivation of the PLC did not change the overall pattern of results for effects sex, stimulus value, or number of offers, as reported above during the choice learning sessions. As such, sex as a biological variable was not assessed for effects of prelimbic inactivation. Choice preference did not change at any dose of muscimol (p=0.122, F(3,48)=2.029) (Figure 6A); however, response latencies were significantly altered by inactivation of the PLC. With increasing concentrations of muscimol, both males and females had shorter response latencies (p=8.49e-08, F(3,329)=12.823). Post hoc testing found pairwise differences between the highest concentration of muscimol and PBS (p=0.0005) and the lowest concentration of muscimol (p=0.00768) (Figure 6B). ExGauss modeling (not shown) revealed an overall decreases in the Gaussian mean (mu) (p=0.015998, F(3,176)=3.534) and exponential variability (tau) (p=0.0054, F(3,176)=4.367). Post hoc testing of tau found a pairwise difference between PBS and the highest concentration of muscimol (p=0.0086184).

**Figure 5.**
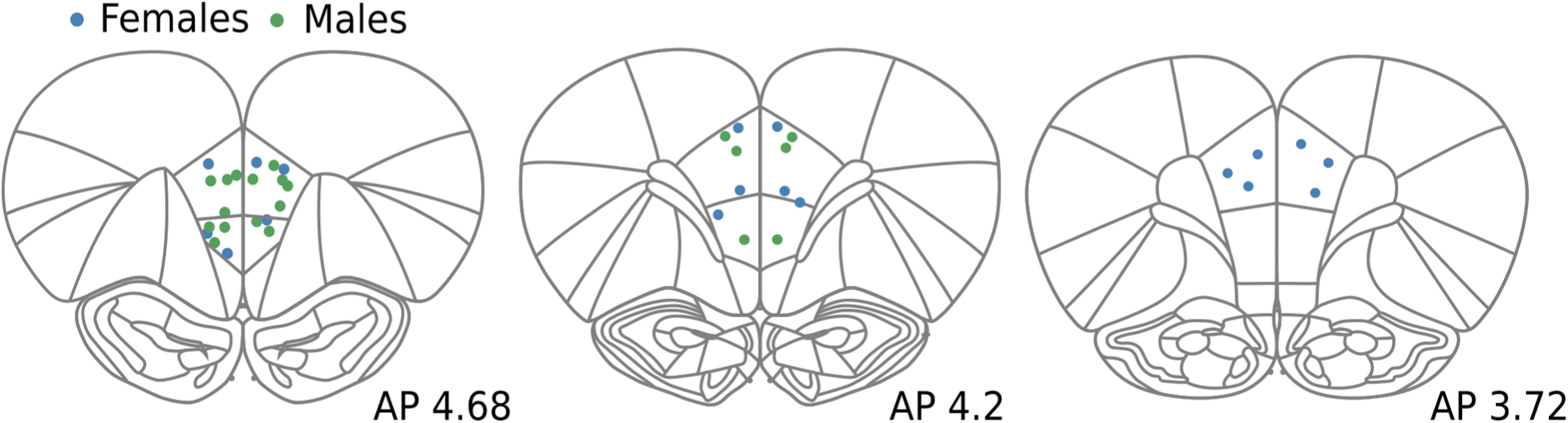
Maps of cannula placements. All cannula tips were localized to the prelimbic cortex (area 32: Paxinos and Watson, 2014) between 3.72 and 4.68 mm anterior to bregma. Cannulas from females are shown as blue points. Cannulas from males are shown as green points.

**Figure 6.**
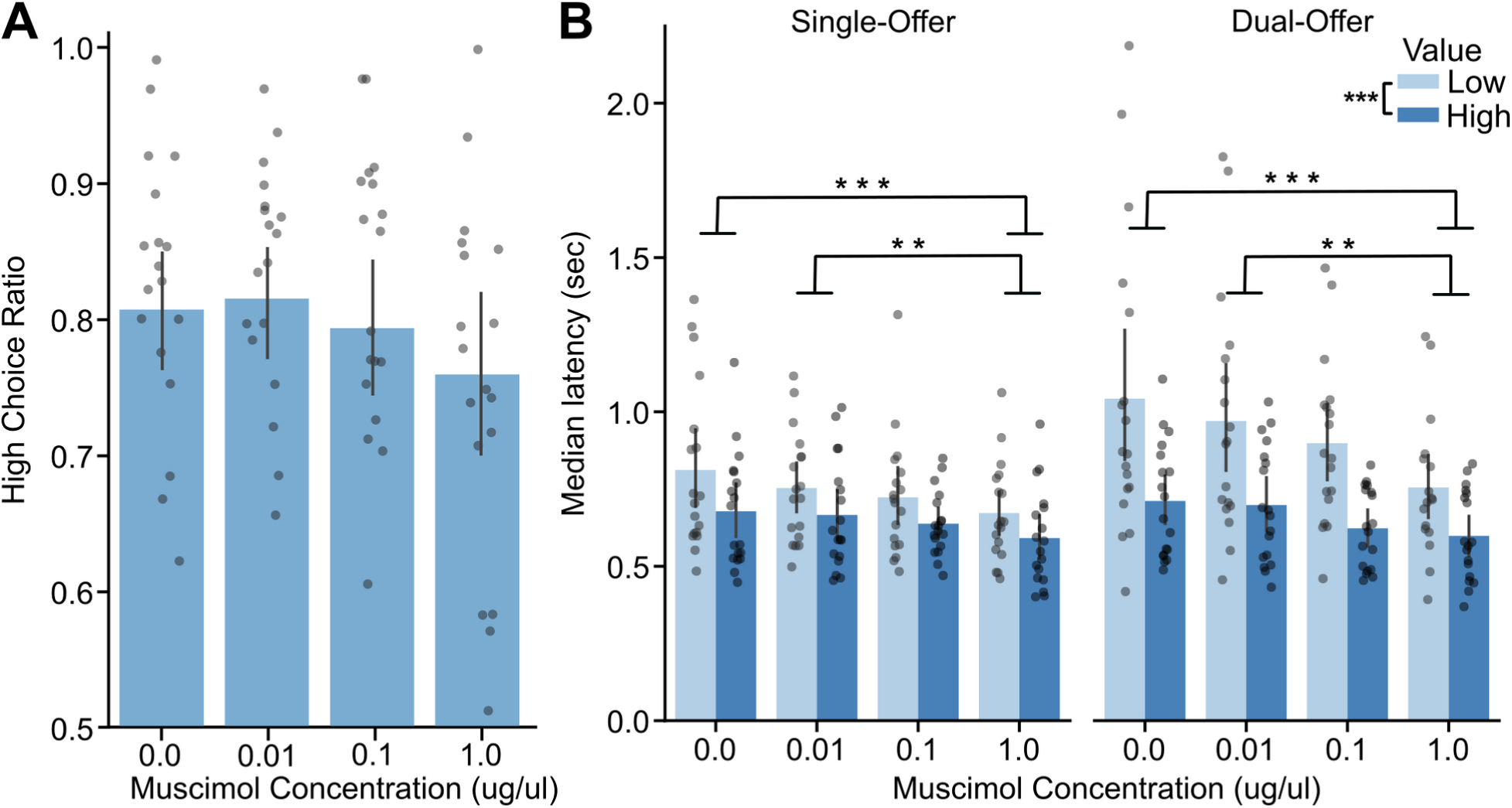
Effects of PLC inactivation on choice preference and latency. **A.** High value preference, the fraction of trials in which the rats chose the higher value stimulus, was not affected by inactivation of PLC. **B.** Median response latency reduced by PLC inactivation in males and females at the highest dose of muscimol (1 µg/µl) compared to the control session and the lowest dose of muscimol. These effects were found on single and dual offer trials. Rats retained sensitivity to reward value with PLC inactivated. Asterisks: *** p<0.001, ** p<0.01, * p<0.05.

To understand how inactivation of PLC affects the decision process, we fit HDDM models to the data from the series of muscimol inactivation sessions. Separate models were fit for each of the three DDM parameters, in which one parameter was allowed to freely vary over doses of muscimol and the other parameters were estimated globally. As above for our analyses of choice learning, we noted differences over doses of muscimol in the parameters based on the probability that the distributions of estimated parameter values did not overlap in either their lower or upper limits. The distributions for drift rate and non-decision time fully overlapping over the range of doses for muscimol (Figure 7A,C). Threshold decreased with increasing dose of muscimol (Figure 7B), and the probabilities that threshold was greater in the sessions with PBS and 0.01 µg/µl of muscimol compared to the sessions with 1.0 µg/µl of muscimol were 0.0028 and 0.0072, respectively. These differences are denoted by two asterisks in Figure 7B.

**Figure 7.**
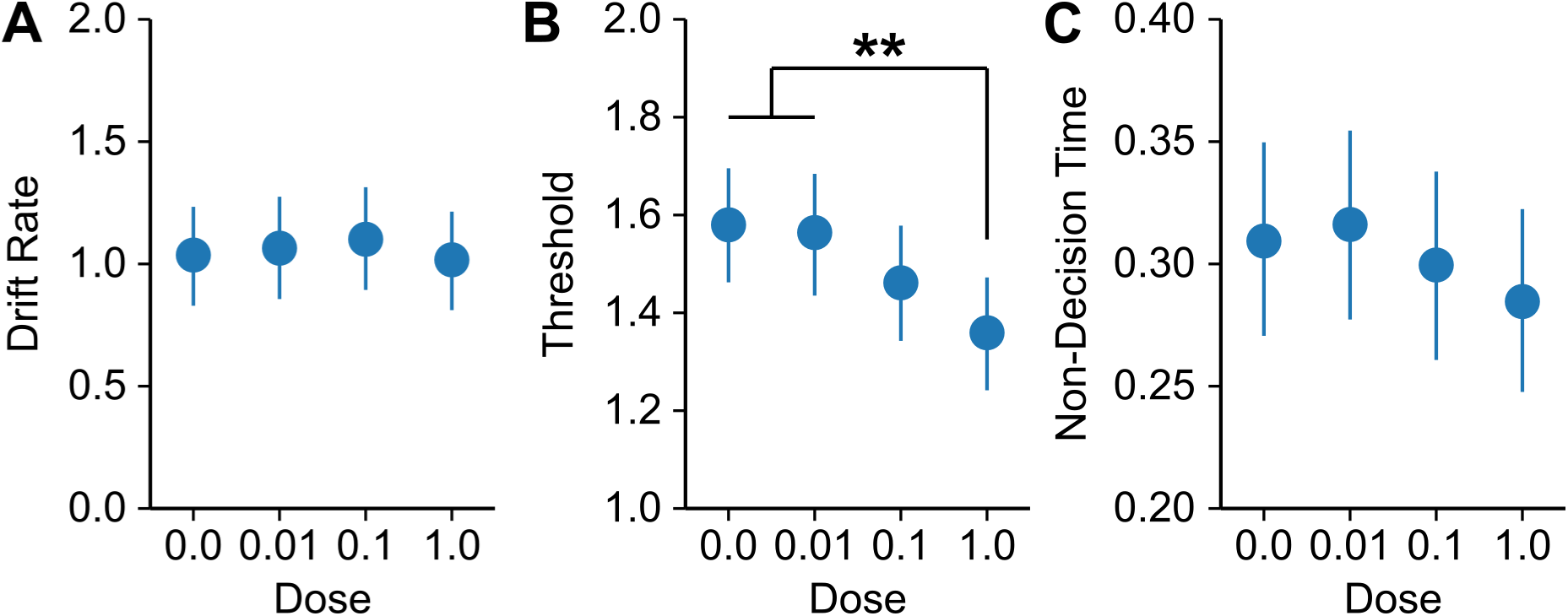
Effects of PLC inactivation on parameters from drift diffusion modeling. A. Drift rate was not affected by PLC inactivation. B. Threshold was reduced by PLC inactivation. Threshold was higher after infusions of vehicle and the lowest dose of muscimol compared to infusions of the highest dose of muscimol. C. Non-decision time was not affected by PLC inactivation. Asterisks: *** p<0.001, ** p<0.01, * p<0.05.

Other behavioral measures were evaluated and found to show effects of PLC inactivation. For reward retrieval, a significant interaction between muscimol concentration and sex (p=0.019477, F(3,239)=3.361) revealed that only females had shorter reward retrieval with increasing concentrations. Post hoc testing in females revealed that the highest concentration resulted in shorter reward retrieval compared to PBS (p=0.0005095) and the lowest concentration (p=0.0377331). There was no effect of muscimol on ITI or time at spout. Finally, there was no side bias caused with increasing muscimol concentrations (p=0.0782, F(3,48)=2.413).

Four males and four females were further assessed for effects of unilateral inactivation. Latencies of ipsilateral and contralateral responses to the hemisphere of the infusion did not differ from one another (p=0.306028, F(1,104)=1.058). There was no side bias caused by unilateral infusions of muscimol (p=0.5379, F(1,6)=0.426), and side bias was weakly negatively correlated to high value preference (r(14)=-0.491, p=0.05558).

### Effects of mu opioid receptor stimulation in the prelimbic cortex

In sessions with DAMGO infusions (1 μg/μl), effects of sex, value, and trial type were the same as those reported during acquisition and were unaffected by DAMGO. As such, sex as a biological variable was not assessed for effects of DAMGO. Similar to reversible inactivation, choice preference was not affected by DAMGO (p=0.622, F(1,16)=0.253) (Figure 8A). However, in contrast to muscimol, DAMGO resulted in longer response latencies (p=5.32e-08, F(1,111)=34.089) (Figure 8B). ExGauss modeling (not shown) found corresponding evidence for increases in the Gaussian mean (mu) under DAMGO (p=0.000123, F(1,80)=16.303) without major effects on exponential variability (tau) (p=0.05507, F(1,80)=3.79). Other behavioral measures included reward retrieval being longer in sessions with DAMGO in PLC (p=6.62e-16, F(1,111)=89.300) and ITI (p=2.42e-05, F(1,111)=19.434) and there was no evidence for a side bias caused by DAMGO (p=0.169, F(1,16)=2.078).

**Figure 8.**
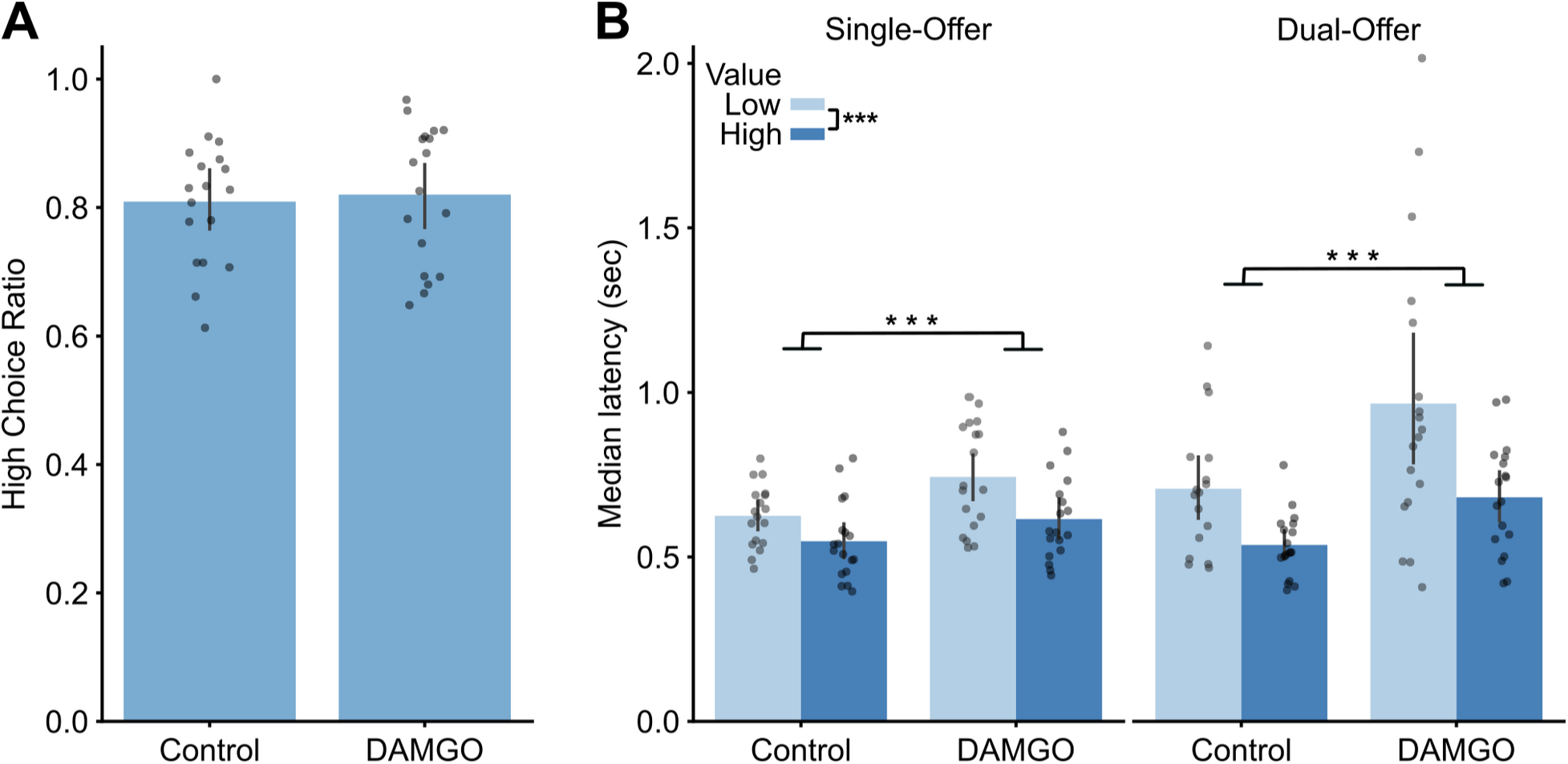
Effects of DAMGO in PLC on choice preference and latency. A. High value preference, the fraction of trials in which the rats chose the higher value stimulus, was not affected by infusions of DAMGO into PLC. B. Median response latency increased following infusions of DAMGO into PLC. This effect was found on single and dual offer trials. Rats retained sensitivity to reward value with DAMGO infused into PLC. Asterisks: *** p<0.001, ** p<0.01, * p<0.05.

DDM modeling revealed non-specific effects of cortical opioid stimulation that contrasted with the effects of PLC inactivation (Figure 9). Drift rate decreased under DAMGO, and the probability that drift rate was lower in the control sessions compared to the drug sessions was 0.0492. (denoted by a single asterisk in Figure 9A). Threshold increased under DAMGO, and the probability that threshold was greater in the control sessions compared to the drug sessions was 0.008 (denoted by double asterisks in Figure 9B). Finally, non-decision time increased under DAMGO, and the probability that non-decision time was greater in the control sessions compared to the drug sessions was 0.0056 (denoted by double asterisks in Figure 9C).

**Figure 9.**
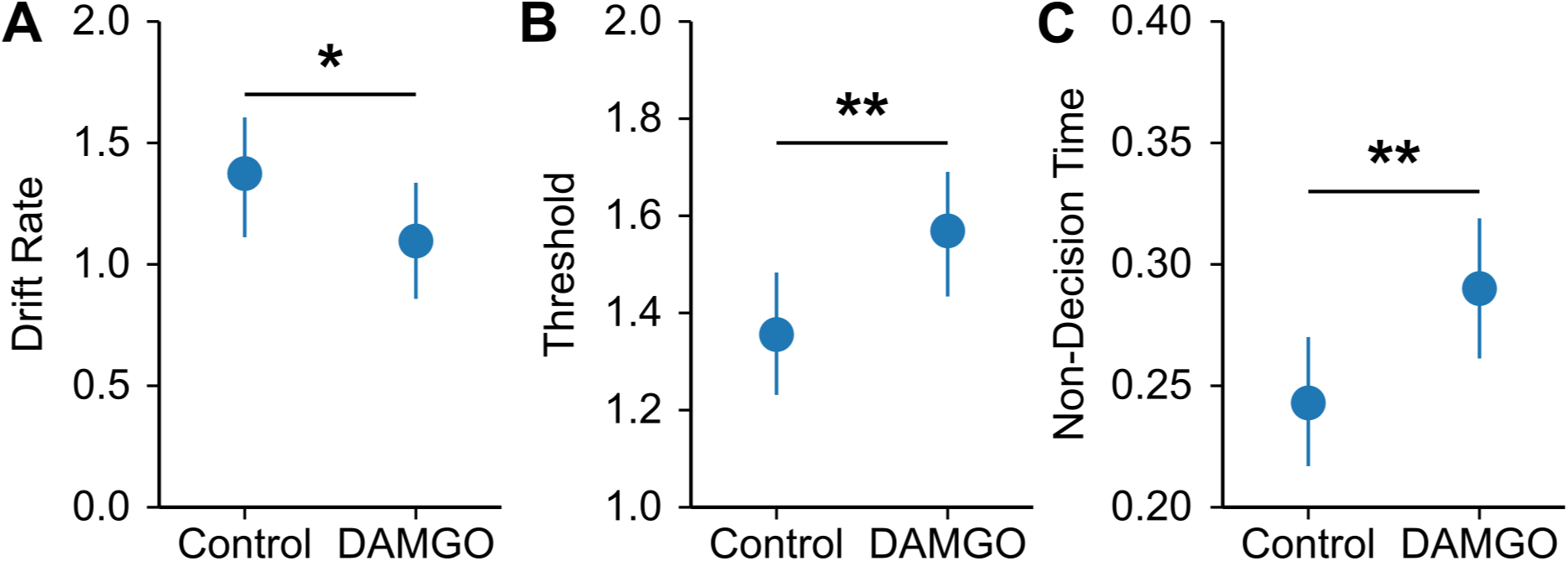
Effects of infusions of DAMGO into PLC on parameters from drift diffusion modeling. A. Drift rate reduced in DAMGO sessions compared to the control sessions. B. Threshold increased in DAMGO sessions compared to the control sessions. C. Non-decision time increased in DAMGO sessions compared to the control sessions. Asterisks: *** p<0.001, ** p<0.01, * p<0.05.

## Discussion

We explored how the prelimbic cortex (PLC) influences decision-making. Our study was inspired by a previous human brain imaging study (Domenech & Dreher, 2010) that suggested that the anterior cingulate cortex, which includes the prelimbic cortex, tracks the amount of information needed to make a choice, i.e. the decision threshold. We used intra-cortical pharmacology to temporarily alter activity in the PLC of rats. This involved either temporarily inactivating the PLC using a the GABA-A agonist muscimol or stimulating PLC using the mu-opioid agonist DAMGO. Our results showed that changing PLC activity impacted how quickly, but not how accurately, rats made decisions. Interestingly, these effects mirrored changes observed in the human study, suggesting a causal role of the PLC in maintaining the decision threshold. Our findings suggest the PLC acts like a “brake” on decision-making, allowing rats to process more information before making a choice. Along the way, we observed a differences between male and female rats in how they learned to make choices. Female rats made faster decisions based on what they learned about the values of the task stimuli and maintained consistent performance during the period of choice learning. In contrast, male rats initially made slower decisions but improved over time, and also showed decreases in their decision thresholds. Overall, our research provides direct evidence that the PLC plays a crucial role in decision-making by regulating the amount of information needed before a choice is made. Further studies are needed to explore the observed sex differences found in our study and understand how individual neurons within the PLC might change during the initial learning phase of decision making tasks.

### A potential sex difference in the learning and performance of decision making tasks

During single-offer instrumental training, all of the animals reduced latencies over sessions, and females had shorter latencies than males (Figure 2A). During dual-offer acquisition, males became faster with experience, while females remained consistent across sessions (Figure 2B). Importantly, females had shorter latencies than males specifically when choosing the high value stimulus and consistently chose the high value stimulus more than males across sessions (Figure 2B,C). This suggests that the females responded more selectively and faster to the high value stimulus.

It is possible that females may have developed a stronger preference to the high value stimulus due to an innate sex difference in consumption behavior, where females have been shown to have a stronger preference for sucrose (Sclafani et al., 1987; Reichelt et al., 2016). All animals demonstrated slower latencies for dual-offer trials (Figure 2C). This difference based on trial type was observed only in latencies and not in other behavioral measures (reward retrieval, ITI, etc.), suggesting a comparative process while making a choice. The effect of value (responding faster for the high value stimulus) was also only observed in latencies, suggesting that value only impacts the speed of the choice itself.

It would be expected that on dual-offer trials, animals would choose high value four times as much as the low value, a phenomenon known as ‘matching’ behavior (Herrnstein, 1961). Females demonstrated matching, choosing the high value stimulus for approximately 80% of dual-offer trials, while males chose high 70-75% of the time. Choosing the more rewarding option less than expected is known as “undermatching”, and is found more commonly than perfect matching (Bari and Gershman, 2023; Baum, 1974; Baum, 1979; Houston et al., 2021; Trepka et al., 2021). In the present study, it is possible that males undermatch due to “policy complexity”, which is the assumption that a decision is made while simultaneously attempting to maximize reward while minimizing cognitive cost (Bari and Gershman, 2023). Alternatively, males may undermatch because the cognitive cost of the decision is low since either outcome (4% or 16% sucrose) is rewarding. Females, perhaps because of their innate sucrose preference, might have demonstrated matching because 16% sucrose is distinctly more sapid and rewarding. This interpretation follows the well-established tendency of female rats to consume proportionally more carbohydrates than males rats (Sclafani et al., 1987).

Our computational modeling suggests that females had faster integration of the task stimuli and faster sensorimotor preparation (higher drift rates, briefer non-decision times), allowing them to make choices more quickly than males. Response latencies of females were less variable (lower tau from ExGauss modeling) when they responded for the high value stimulus, a difference not observed in males (Figure 3B). These findings may explain why females responded faster than males specifically for the high value stimulus (Figure 2). Finally, the variability of response latencies decreased with learning in males (Figure 3B), likely explaining the speeding of latencies of males over sessions (Figure 2C).

### Inactivation of PLC speeds performance without affecting choice

With inactivation, all animals responded faster with no impact on choice preference (Figure 6). Complementary drift diffusion modeling revealed that the decision threshold, but not drift rate or non-decision time, decreased with PLC inactivated (Figure 7B). This finding suggests that, with PLC inactivated, the rats made decisions based on less evidence. We suggest that this finding might follow from a reduction in behavioral disinhibition, a process that has been shown to depend on PLC in many published studies (see Bari and Robbins, 2013 for review).

There was no impact of PLC inactivation on the animals’ choice preferences and the inactivations did not induce spatial bias. The observed effects on response latency without affecting choice preference is consistent with past studies which show that PLC mediates inhibitory control. For example, lesions or inactivation of PLC increase perseverative responding (Chudasama & Muir, 2001), increase stop-signal reaction times (Bari et al., 2011), and decrease conditioned suppression of sucrose seeking (Limpens et al., 2015). These effects together suggest a general role of PLC in inhibitory control, and not inhibition of a specific actions. The same general effects of PLC inactivation were found in the present study. Rats showed reduced response latencies, and these effects were found for both single-offer and dual-offer trials. PLC inactivation speeded performance overall, independent of the needs of the animals to make choices between the task stimuli.

In this study, effects of inactivation were specific to response latencies. It is possible that if a different frontal region were targeted for drug infusions, there may be a choice preference effect. For instance, OFC lesions in the five-choice serial reaction time task resulted in reduced accuracy, which can be interpreted as decreased attention to stimuli (Chudasama et al., 2003). If OFC were targeted in this study, it is possible that a similar lack of attention to stimuli would take place, resulting in altered choice preference. Additionally, animals in this study were instrumentally trained and had experienced dual-offer trials prior to drug infusions. It has been shown that PLC is necessary for the acquisition of goal-directed behaviors, and that training prior to inactivation or lesioning does not impact behaviors with regard to the value of reward (Killcross and Coutureau, 2003; Tran-Tu-Yen et al., 2009). This is likely another indication of why there were no effects on choice preference in the present study. Choice preference may have been affected if PLC were inactivated during task acquisition when animals were learning stimulus-reward contingencies, or when they experienced a dual-offer trial for the first time.

### Mu opioid stimulation slows performance without affecting choice

Studies that stimulate mu opioid receptors in medial parts of the rodent frontal cortex, ventral to the region that was the focus of the present study, have reported opposing effects of inactivation using muscimol and stimulation using infusions of the mu opioid agonist DAMGO (Mena et al., 2011). A recent study by White and Laubach (2022) used this approach to study the role of the prelimbic cortex in a reward-guided consummatory task. They found effects of inactivation of prelimbic cortex, but not opioid receptor stimulation, associated with the rats being able to maintain consumption. To our knowledge, no published study has examined effects of mu opioid stimulation in operant choice tasks, as in the present study.

We found that in all rats infusions of DAMGO resulted in slower responses and no effects on choice preference (Figure 8). Drift diffusion models found effects of DAMGO across all three parameters of the drift diffusion models (Figure 9), indicating that mu opioid stimulation of PLC had non-specific effects on the decision process.

Muscimol inactivates an area by directly increasing GABA tone as a GABA-A agonist, while stimulation of mu opioid receptors with DAMGO has been shown to indirectly inhibit GABA transmission through downstream intracellular cascades after binding on parvalbumin interneurons (Lau et al., 2020). The distinct effects of inactivation and mu opioid stimulation were likely due to distinct effects of the two drugs on neural activity in PLC. Muscimol has well-known effects on neural activity, and has been shown to reduce spiking activity (Martin and Ghez, 1999; van Duuren et al., 2007). By contrast, little is known about how mu opioid stimulation affects brain activity in awake, behaving animals. More work should be done to investigate the role of the opioid system throughout the frontal cortex and its impact on decision making.

## Acknowledgments

We thank Dr. David Kearns and Dr. Yogita Chudasama for their helpful comments. This work was supported by a Pilot Award from the DC Center for AIDS Research and an NIH grant to ML (1R15DA046375-01A1).

## Notes

https://github.com/LaubachLab/PLC-DDM

### Competing Interest Statement

The authors have declared no competing interest.

